# Comparison of EEG source reconstructed functional networks in healthy subjects elicited during visual oddball task

**DOI:** 10.1101/639815

**Authors:** Kang Wei Thee, Humaira Nisar, Kim Ho Yeap, Wei Meng Tan

## Abstract

In this paper we have reconstructed electroencephalography (EEG) sources using weighted Minimum Norm Estimator (wMNE) for visual oddball experiment to estimate brain functional networks. Secondly we have evaluated the impact of time-frequency decomposition algorithms and scout functions on brain functional networks estimation using phase-locked value (PLV). Lastly, we compared the difference between target stimuli with response (TR) and non-target with no response (NTNR) cases in terms of brain functional connectivity (FC). We acquired the EEG data from 20 healthy participants using 129 channels EEG sensor array for visual oddball experiment. Three scout functions: i) MEAN, ii) MAX and iii) PCA were used to extract the regional time series signals. We transformed the regional time series signals into complex form using two methods: i) Wavelet Transform (WT) and ii) Hilbert Transform (HT). The instantaneous phases were extracted from the complex form of the regional time series signals. The FC was estimated using PLV. The joint capacity of the time-frequency decomposition algorithms/scout functions applied to reconstructed EEG sources was evaluated using two criteria: i) localization index (LI) and ii) R. The difference in FC between TR and NTNR cases was evaluated using these two criteria. Our results show that the WT has higher impact on LI values and it is better than HT in terms of consistency of the results as the standard deviation (SD) of WT is lower. In addition, WT/PCA pair is better than other pairs in terms of consistency as the SD of the pair is lower. This pair is able to estimate the connectivity within parietal region which corresponds to P300 response; although WT/MEAN is also able to do that, However, WT/PCA has lower SD than WT/MEAN. Lastly, the differences in connectivity between TR and NTNR cases over parietal, central, right temporal and limbic regions which correspond to target detection, P300 response and motor response were observed. Therefore, we conclude that the output of the connectivity estimation might be affected by time-frequency decomposition algorithms/scout functions pairs. Among the pairs, WT/PCA yields best results for the visual oddball task. Moreover, TR and NTNR cases are different in terms of estimated functional networks.

## Introduction

Brain connectivity may be defined as the links between different units of the brain. These links can be anatomical or structural referred to as structural connectivity (SC), statistical dependencies known as functional connectivity (FC) and due to causal interaction known as effective connectivity (EC).

Neuroimaging techniques are widely used to estimate the brain cortical networks involved in the normal brain cognitive functions as well as in neurological diseases [1-6]. Diffusion Tensor Imaging (DTI) can be adapted to estimate SC with higher capability [7-9]. However, this technique is unable to estimate the dynamic connectivity among the cortical regions. The functional Magnetic Resonance Imaging (fMRI) technique is widely utilized to characterize the cortical FC [10-12] as it provides excellent spatial resolution. However fMRI has relatively lower temporal resolution [13,14] (slow sampling rate; ∼1s). Hence it cannot capture the dynamic activity of the cognitive processes that have extremely short duration. These processes require excellent temporal resolution to capture the dynamic changes. Thus, Electroencephalography (EEG) can solve this problem. EEG is used to measure the scalp electrical potentials using sensor-array. EEG provides excellent temporal resolution. Hence, EEG data with suitable signal processing techniques can provide relative information regarding brain FC elicited during cognitive activities [15-18]. However, EEG suffers from low spatial resolution issue due to volume conduction [19,20]. Same underlying source may influence the EEG signals acquired from two neighbourhood sensors [21]. Thus, estimation of connectivity on EEG signals measured from scalp does not exactly convey the true neural linkages among two brain areas [22]. Some approaches have been proposed to solve this issue. For example, non-linear methods like phase-locked value (PLV) [23] and imaginary coherence (IC) [24] have been proposed for FC estimation as these approaches are insensitive to volume conduction. The application of these methods on reconstructed EEG source signals provides superior spatial and temporal resolution [25-27].

Many algorithms have been proposed for EEG source reconstruction [28-31]. For FC estimation, several linear and non-linear methods have been developed [27,32,33]. In [34-37], some studies have discussed the application of the connectivity estimation methods on the dynamics source signals reconstructed from scalp EEG. These methods provide high efficiency for the FC estimation as the FC is directly estimated from the source space (cortex level). In this context firstly the algorithms used to perform EEG source localization by solving the ill-posed EEG inverse problem for source localization have to be implemented, followed by the FC estimation in the source space.

Several approaches have been proposed in the past decades in order to solve the ill-posed EEG inverse problem. For example, Minimum Norm Estimator (MNE) [38], Depth-weighted Minimum L2 Norm Estimator (wMNE) [39], Low Resolution Brain Electromagnetic Tomography (LORETA) [40], standardized Low Resolution Brain Electromagnetic Tomography (sLORETA) [41] and others. These methods are widely used to solve the ill-posed inverse problem for EEG source localization in recent researches [42-45]. As an outcome, the spatial resolution of the EEG has been improved using EEG source localization.

Secondly the brain FC estimation methods are categorized into linear and non-linear methods. The linear approaches include cross-correlation [46] and coherence [47], whereas non-linear methods include mutual information [48], phase synchronization [49] etc. Linear and non-linear methods are widely used for FC estimations in the sensor space [50-52] as well as source space [53-56].

Based on the literature review it is observed that the estimated FC depends on the algorithms to solve the EEG ill-posed inverse problem and the methods for connectivity estimation. Hassan et al. reported that the use of wMNE in conjunction with the phase-locking value (PLV) provides better results as compared to the other combinations in the sensor space [26,57]. This combination has been adapted for FC analysis in the source space [25,58-60].

In this research, we planned to utilized wMNE to reconstruct the dipolar sources from EEG by solving the ill-posed inverse problem. Then, we applied PLV to estimate the pairwise connections between the regions-of-interest (ROIs). Time-frequency decomposition algorithms like complex Morlet Wavelet Transform (WT) and Hilbert Transform (HT) [33,61] can be adapted to transform the signals in time domain into complex time-frequency domain for instantaneous phase extraction. These two different time-frequency decomposition algorithms may have an effect on the extraction of instantaneous phases that may affect the FC estimation using PLV. In our study, we applied PLV on 148 regional time series signals to estimate the FC.

The estimated dipolar sources on the cortex were downsampled into 148 regions based on Destriuex atlas [62] to form regional time series. The dipolar sources within a region were grouped together into one unique source. This unique source is then used to express the cortical activity of that particular region from Destriuex atlas. Brainstorm toolbox [63] provides few options (scout functions) to perform the grouping. The scount functions are MEAN, PCA, MAX and others. MEAN averages all dipolar sources within a cortical region to produce one unique source for that particular cortical region. Using MAX, the unique source selected from the maximum sources across all the dipolar sources within that particular cortical region. And, PCA takes the first mode of the PCA decomposition of all the sources within a cortical region to form a unique source of that particular region. Different regional time series can be generated using different scout functions. Thus, the scout functions can affect the extraction of instantaneous phases from regional time series signals which may affect the FC estimation using PLV.

Hence we believe that the network differences between two oddball cases are significant. In addition we also assume that different time-frequency decomposition algorithms and scout functions may have slightly different impact on FC estimation.

Based on our hypotheses, we have two main objectives. 1) To evaluate the differences in terms of functional connectivity among oddball cases in the source space. Previously the evaluation was done in the sensor space [50,51,64], 2) To assess the joint capacity of the time-frequency decomposition algorithms / scout functions applied to reconstructed EEG sources to evaluate brain FC elicited by our oddball task; as the joint capacity has not been evaluated by other studies. In this study, we also assess the impact of the scout functions and time-frequency decomposition algorithms on FC estimation.

## Methodology

The flow chart of the research is depicted in Fig 1. Dense EEG electrode arrays was used to acquire the EEG data for visual oddball experiment. Secondly the dipolar sources were reconstructed from clean scalp EEG data using wMNE. Regional time series was computed based on Destrieux atlas using different down-sampling scout functions (PCA, MAX and MEAN). In the third step, time-frequency decomposition was carried out using Morlet WT and HT. After decomposition, regional time series in complex form in gamma band was used to extract the instantaneous phases. The extracted instantaneous phases were used to estimate the FC using PLV. Proportional thresholding was applied to retain only 10% of strongest connectivity. Lastly, performance of each combination of the time-frequency decomposition algorithms with scout functions was evaluated based on the Localization index (LI) and R criterions. Statistical tests were performed to evaluate the level of significance.

**Fig 1.**
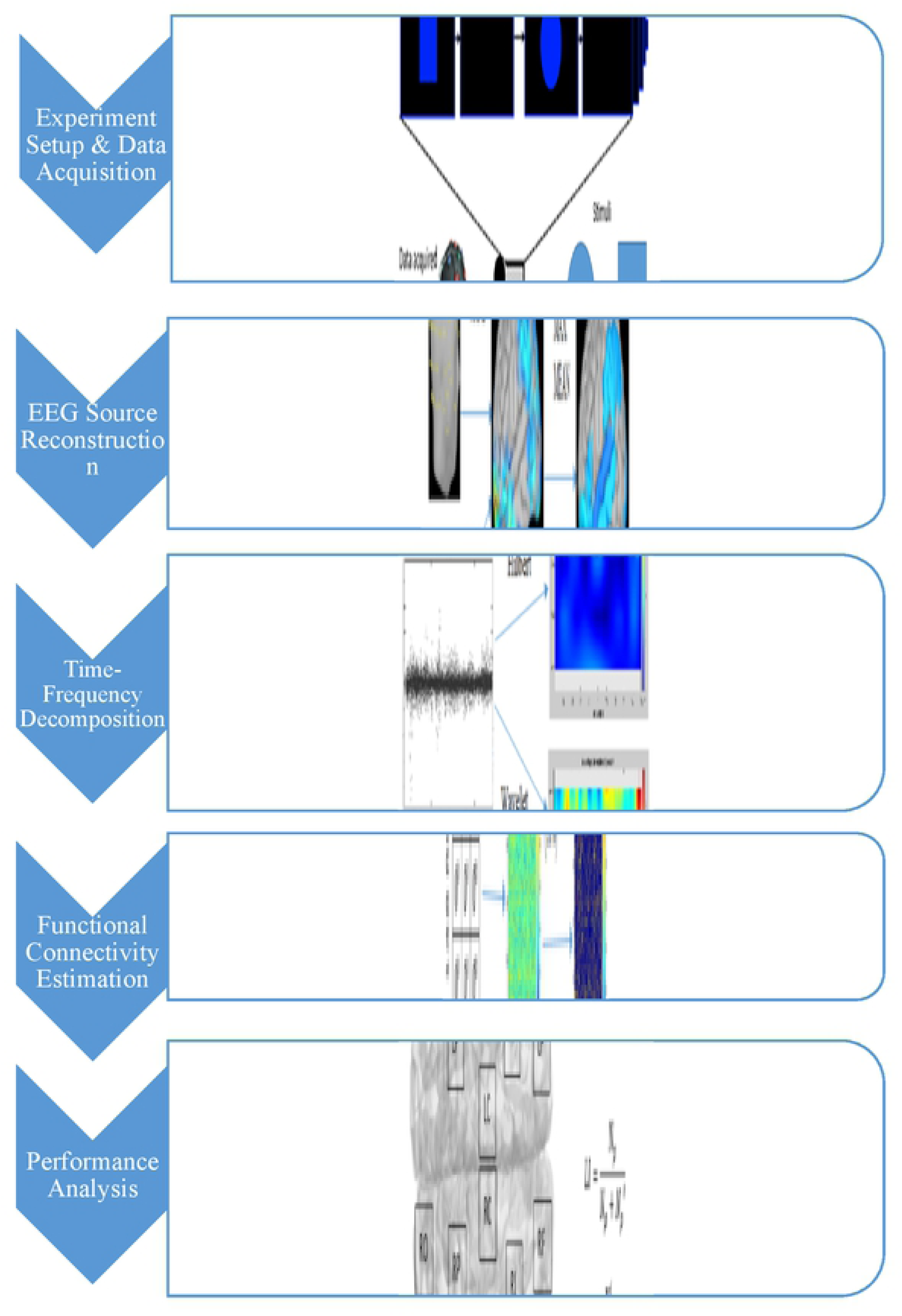
Flow chart of the study.

### Experiment setup and data acquisition

In an oddball paradigm, the researchers asked the participants to distinguish the novel stimuli (target) within a series of randomly displayed frequent stimuli (non-target).

We randomly presented the target stimuli (circle) and non-target (square) stimuli on the computer monitor for 500ms during our visual oddball paradigm [64]. The fixation time was set as 1000ms. During the fixation, an empty dark screen was presented. We requested the subjects to pay attention towards the monitor. They have to make motor response by pressing the keyboard button when the target stimuli appears on the computer monitor. When the the non-target stimuli appears; the motor response is not required. Total 135 visual stimuli were presented on the monitor. 40 out of 135 stimuli were the target stimuli, whereas 95 stimuli were the non-target stimuli. The stimuli were projected on the monitor randomly.

Our oddball paradigm is categorized into 4 different oddball cases. The ‘correct’ cases are target stimuli with response (TR) and non-target stimuli with no response (NTNR). While, the ‘incorrect’ cases are target stimuli with no response (TNR) and non-target stimuli with response (NTR). In TR case, the participants correctly respond to the target stimuli, whereas in TNR case the subjects fail to respond to the target stimuli. In NTR case, the subjects respond incorrectly to the non-target stimuli. In NTNR case, the participants did not provide the motor response when non-target stimuli appeared. In this study, we used TR case for the evaluation of the scout functions and time-frequency decomposition algorithms. Moreover, we used TR and NTNR to compare the differences between the two cases in term of connectivity as these two cases are opposite to each other.

The 128-channel sensor array (HydroCel Geodesic Sensor Net) from EGI company with a sampling frequency of 250 Hz was used to acquire EEG data. The EEG data was acquired from 20 right-handed healthy participants with an age of around 19–23 years with normal or corrected-to-normal vision. None of them had a history of substance abuse and a personal or family history of psychiatric or neurological diseases.

The sampling frequency of the EEG acquisition system is 250 Hz. The maximum time period of the epoch is about 500ms. The data acquisition for TR case will be stopped once the subjects provided the motor response, hence the period of the epoch for this case could be less than 500 ms. The raw data was converted into Matlab format by netstation. The high frequency artifacts and DC components were removed using a finite impulse response (FIR) digital filter with a band-pass frequency range from 0.5 Hz to 70 Hz. The EEGLAB function called eegplot() was used to plot the filtered EEG data. The EEG samples that consist of artefacts induced by muscles contraction and eye blinking were manually rejected. For TR case, the clean EEG data were shifted to the right to align with the event when the button is pressed by the participants. For NTNR case, shifting is not required. The data of 2 participants was rejected. For TR and NTNR cases, three trials have been randomly selected from each subject (3 × 18 = 54 trials) to perform the analysis. We performed the analysis 3 times by selecting different trials from each subject for both cases.

### EEG source reconstruction

The generative model of EEG data, *E(t)* can be expressed as linear consolidation of time-varying current dipole sources *S(t)* with 3-dimensional (3D) orientation:

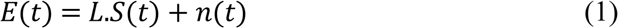

where *n(t)* denotes the additive noise and *L* denotes the lead fields matrix of the dipolar sources [26]. This process is essentially known as forward modelling. On the other hand, inverse modelling is known as the process of solving the inverse problem by reversing the step of estimating *S(t)* given *E(t)* and *L*.

#### Forward modelling

The physical process of the neuronal current propagation from brain cortical surfaces to the EEG electrodes on the scalp is described by the lead field *L*. The electrical conductivities and geometry of the tissues in the head are required to compute *L*.

Ideally, the geometrical model of an individual’s head should be obtained from various structural MRI and digitized sensor positions. Unfortunately, taking individual MRI is costly. Thus, anatomical templates are commonly used in EEG source analysis. We utilized ICBM152 template (a non-linear average of the MRI images of 152 individual’s heads) for our study [65,66]. To obtain the electrical properties of the tissues in the head, we utilized boundary element method (BEM) [67]. The realistically-shaped shells that represent the brain, scalp and skull are included in BEM.

Based on Destriuex atlas, the cortical surface was partitioned into 148 ROIs. EEG forward modelling was done using Brainstorm toolbox. The surface meshes of the scalp, skull and brain which are realistically-shaped were extracted from the ICBM152 template. BEM was used to calculate the forward head model (lead field) from these 148 ROIs to the 129 EEG channels as implemented in Open MEEG software package [68] included in Brainstorm toolbox. In this case, we set the number of vertices per layer as 1922 (default). The electrical conductivity for brain, skull and skin were set to 1 S/m, 0.0125 S/m and 1 S/m respectively.

#### Inverse modelling

In inverse modelling, dipolar sources are estimated over the cortical regions given the EEG signals and lead field matrices. MNE [38] imposes L2-norm constraints on the source distribution are efficient as the method is linear in the sensor data. L2-norm is introduced to regularize the problem. A weighted matrix is introduced to MNE algorithm to improve the source estimation in terms of surface sources. This method is known as wMNE [39]. The weighting matrix adjusts the properties of the solution by reducing the bias inherent to MNE algorithm. The source estimated by wMNE is expressed as

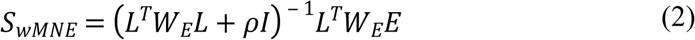

where W_E_ is the diagonal weighting matrix which consists of weighted factor for depth normalization, *ρ* denotes the regularization parameter and *I* is the identity matrix. The inverse estimation of sources was performed by Brainstorm toolbox. The value of *ρ* was set between 0.1 and 0.3 [60]. After that, the dipolar sources were projected on the 3D cortical surface. The regional time series was computed based on three different scout functions offered by the Brainstorm toolbox.

### Time-frequency decomposition

We used HT and WT to decompose the regional time series into time-frequency representation.

#### Hilbert Transform

The regional time series signals were decomposed into gamma band (30 – 60 Hz) using FIR band pass filter. After filtering, HT was applied. HT of a function *f(t)* can be defined as convolution of *f(t)* with 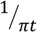. HT of a regional time series signal *x(t)* can be mathematically represented as:

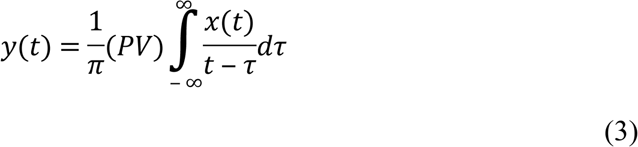

where *PV* denotes the Cauchy principle value [69].

#### Wavelet Transform

WT outputs a time-frequency plane for regional time series signal *x(t)*. During spectral analysis, complex Morlet Wavelet function provides magnitude and phase information of the time series signals [70]. Mathematically, the Wavelet function is defined as follow:

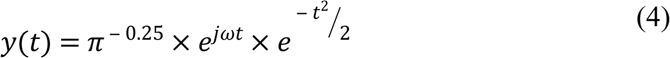

where *y(t)* is a sinusoid in complex form of *e*^*jwt*^ multiplied by a normalization factor (*π ^™ 0.25^*) and Gaussian envelope 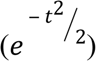. This process is to ensure the Morlet Wavelet has unit energy.

### Functional connectivity estimation

HT and WT convert the time series signal *x(t)* into complex function of time *δ(t)* defined as:

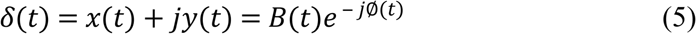

Where ∅*(t)* denotes the instantaneous phase with respect to time and *B(t)* denotes the instantaneous amplitude with respect to time [69]. A MATLAB function *angle()* is used to extract the instantaneous phases of the regional time series signals in radian form.

The phase differences *θ*_*ab*_(*t,n*) between two regional time series signals a and b at time bins *t* and trial *n* were computed as follows:

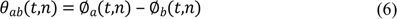

where ∅_*a(t,n)*_ and ∅_*b(t,n)*_ are the instantaneous phase of regional time series signals a and b [71].

An index known as PLV was used to define the degree of synchronization between the two estimated instantaneous phases [23]. Mathematically, PLV is expressed as:

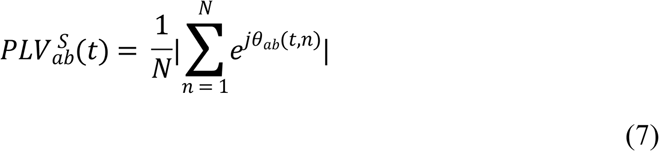

where *S* denotes the subjects and *N* denotes the total number of trials. The grand average of PLV over 18 subjects 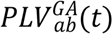 is defined as

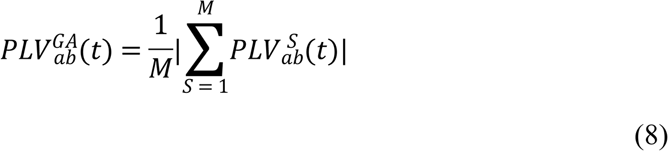

where *M* denotes the total number of subjects. Adjacency matrix was formed to represent the connectivity graph. The connectivity graph was normalized with the 200 ms pre-stimulus baseline using Z-score normalization procedure. The normalized graph is then defined as

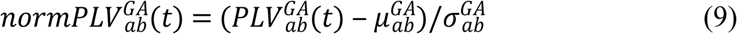

where 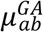 and 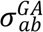 are the grand-averaged mean and standard deviation calculated from the 200 ms pre-stimulus baseline.

Next, the connectivity graph was thresholded using proportional thresholding approach [6,72]. A small percentage (10%) of strongest estimated connections were retained.

### Performance analysis

In Destriuex atlas [62], 148 cortical regions were group into 14 macro ROIs. We predefined the 11 distinct ROIs reported to be implicated in the visual oddball paradigm based on the previously published functional imaging studies of visual oddball task [3,73-89]. The 11 predefined ROIs are listed in bottom of Fig 1.

We performed the performance analysis using two criterions. These criterions quantify the identified networks distributed within the predefined ROIs as ‘correct’ networks whereas the identified networks distributed outside the predefined ROIs are identified as ‘incorrect’ networks.

The first criteria is Localization Index (LI) [26]. LI is a ‘global’ indicator to quantify the performance of the scout functions/time-frequency decomposition algorithms pair. It was computed over the 11 predefined ROIs. LI is defined as the ratio between the number of identified edges within all predefined ROIs and the total number of estimated edges. Mathematically, LI is defined as

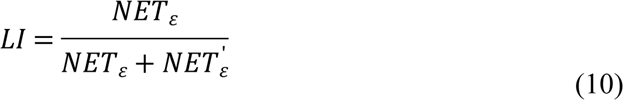

where *NET*_ε_ denotes the number of estimated connections within the 11 predefined ROIs and *NET*^′^_ε_ denotes the quantity of estimated edges outside the predefined ROIs. The LI ranges between 0 (no connections are identified with the predefined ROIs) and 1 (all connections are identified within the predefined ROIs).

Another criteria used for performance analysis is R [26], known as the ‘local’ indicator to quantify the local distribution of identified networks within each ROI. Ratio between quantity of estimated connections within each predefined ROI and total identified connections within all predefined ROIs. Mathematically, the R can be expressed as:

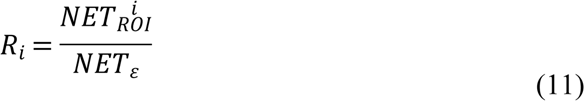

where *i* denotes the 11 predefined ROIs (see Fig 1) and 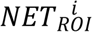 denotes the number of estimated connections within the ROI *i*. The *Ri* criteria ranges from 0 (no estimated edges are found inside ROI *i*) to 1 (all estimated edges are found inside ROI *i*).

## Results and discussion

In this study, we have performed source localization of EEG data acquired during visual oddball task. We used wMNE algorithm to solve the EEG inverse problem. Fig 2A depicts the estimated current density sources on 3-dimensional (3D) cortex between 240 – 500 ms of the epoch. From the top view of the cortex, it is observed that the majority of current sources are intensively distributed on the parietal and central regions. These activations are generally elicited by P300 and motor response. The bottom, left and right views show stronger current sources over the left temporal region. On the front view, peak current sources are mainly distributed on the frontal region. On the back view, lower intensity of current sources are observed over occipital region.

**Fig 2.**
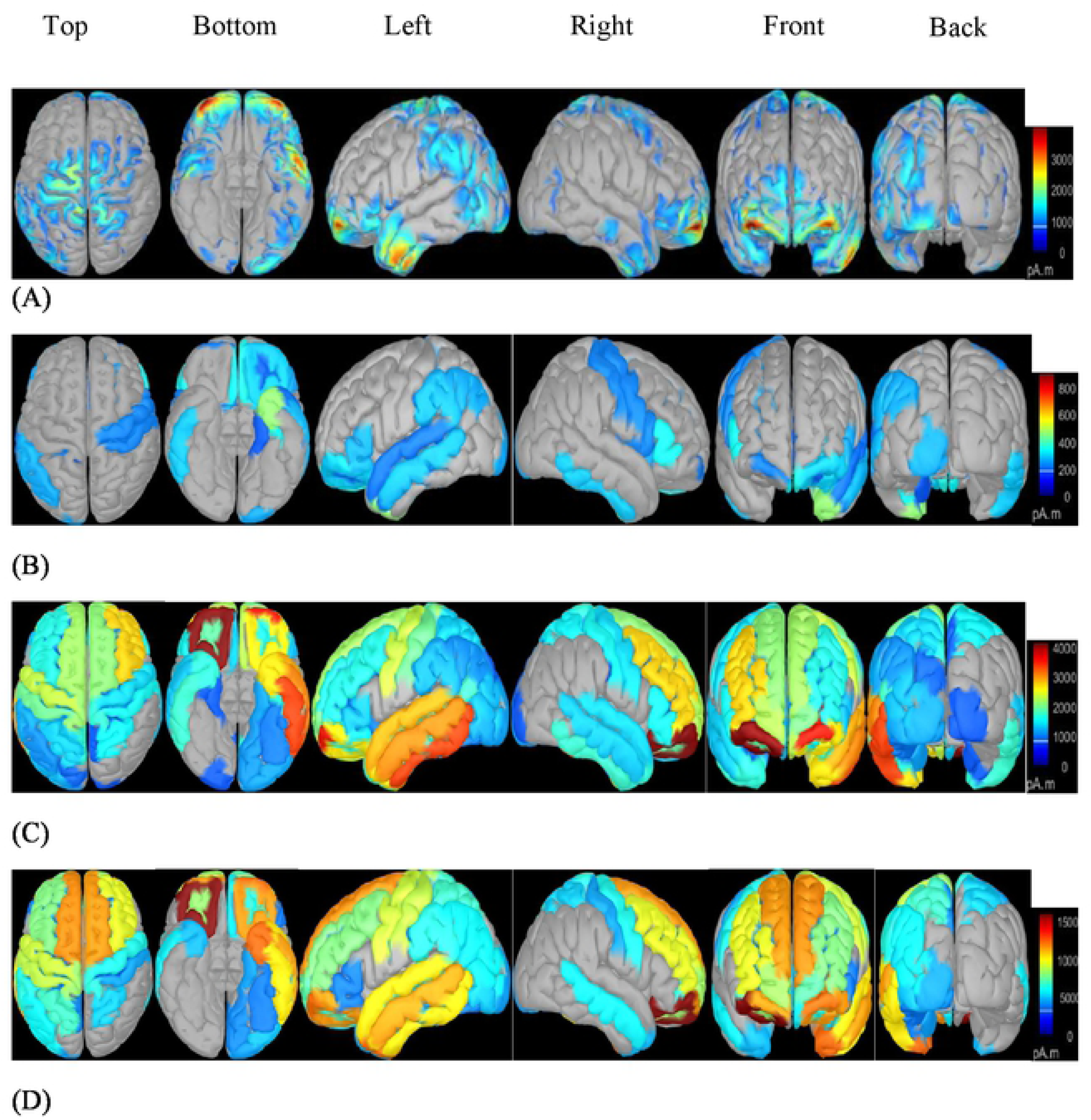
Different views of reconstructed sources on the cortical surface (A) and respective regional cortical sources using different downsamling scout functions: MEAN **(B)**, M.AX (C) and PCA (D).

We partitioned the cortical surface of the brain into 148 regions based on Destrieux cortical atlas. These regions are equally parcelled over right and left hemispheres of the brain surface. These regions are also known as scouts in Brainstorm jargon. We applied several scout functions, i.e. MEAN, MAX and PCA, as source down-sampling algorithm to create the scout (regional) time series (148 regions from Destrieux atlas). Fig 2B - 2D depicts the regional time series projected on 3D cortex between 240 – 500 ms. As we can see on the figure, the regional times series generated by different scout functions are different in terms of current density distribution and intensity. Therefore, scout functions could be a factor that affecting the performance of FC estimation.

We applied WT and HT to decompose the regional time series signals into time-frequency domain. Fig 3 depicts the time-frequency representations of a regional time series (right central region). The peak magnitude in gamma band (30 – 60 Hz) was observed over both spectrograms which is elicited by motor planning and integration. However, both algorithms show different intensity in gamma band as well as in other bands. Therefore, time-frequency decomposition algorithms could be another factor that affects the performance of FC estimation.

**Fig 3.**
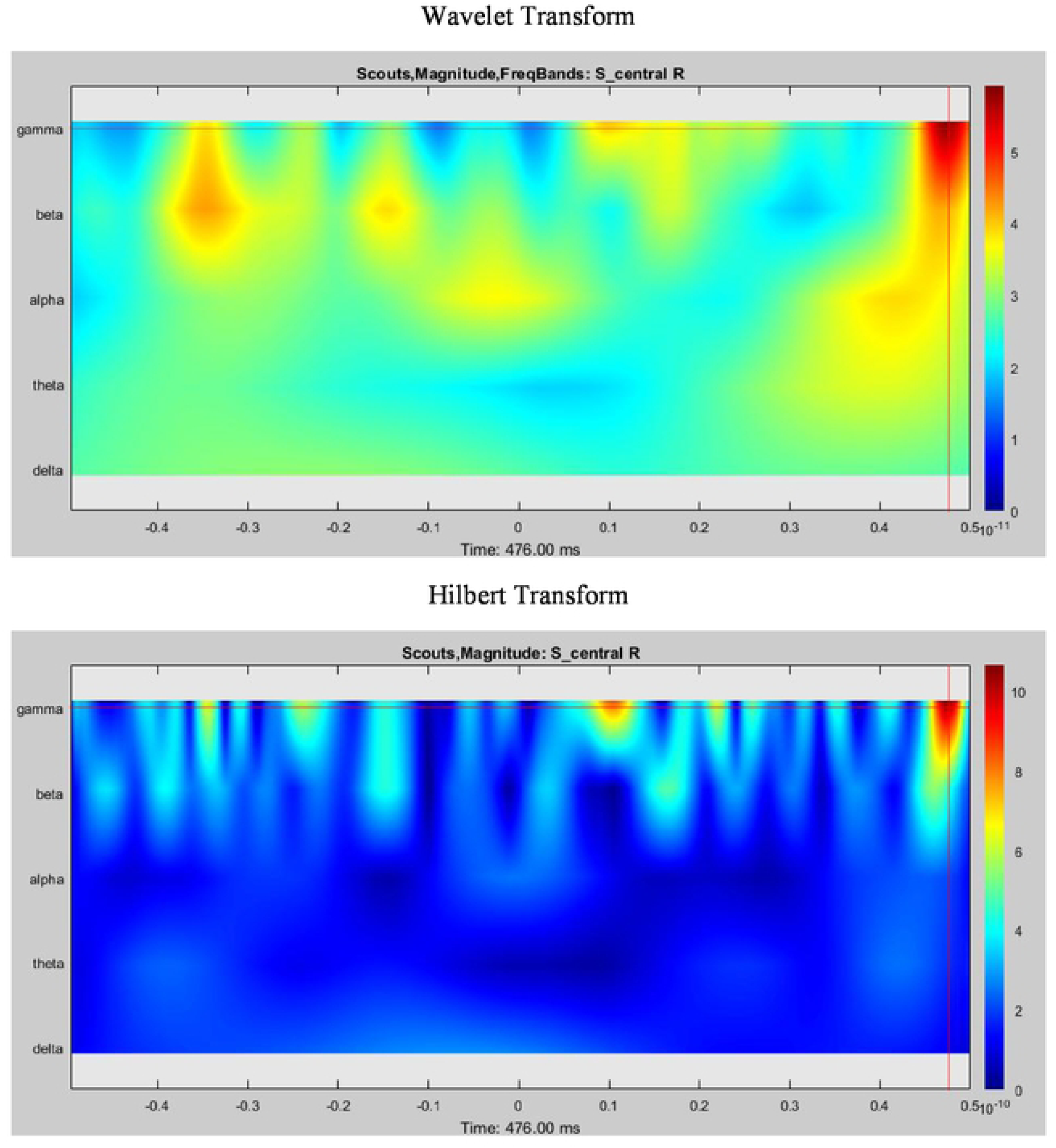
Magnitute of time-freququency decomposition for a central region from Destrieux atlas. Upper panel: Wavelet Transform; Lower panel: Hilbert Transform

We extracted the instantaneous phase of the regional time series in gamma band using PLV to estimate the FC. The connectivity graphs obtained for the 6 different combinations of the time-frequency decomposition algorithms and scout functions are presented in Fig 4. The colour-coded circles are the nodes of the network while the lines linking the two nodes are denoted as an edge. The colours of the nodes show the intensity level of the node degree. The main purpose of the node degree here is to use as a comparative parameter to distinguish the identified networks estimated by different combinations of methodologies. Indeed, the differences can be observed based on the edges. For qualitative visual inspection of the estimated networks, the node degree is compulsory for the comparisons between the estimated networks. The qualitative visual inspection of the estimated edges indicate that the results are highly dependent on the two factors, i.e. time-frequency decomposition algorithms and scout functions.

**Fig 4.**
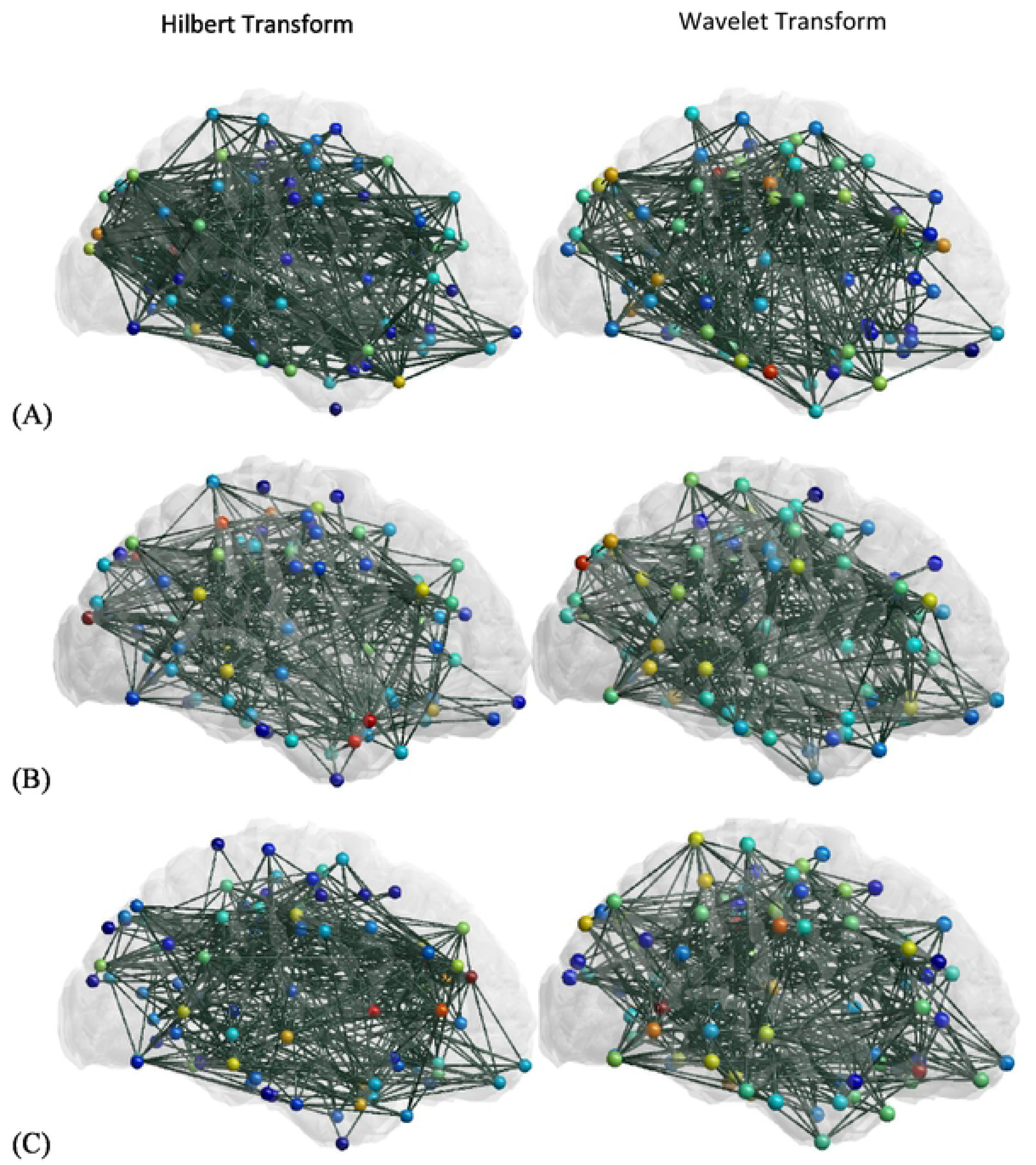
Connectivity estimated by PLV based on the instantaneous phases extracted using Hilbert Transform (left panel) and Wavelet Transform (right panel) from regional time series downsampled by different scout functions: **(A).** MEAN; **(B).** MAX; (C). PCA.

The performance analysis was carried out on TR case using **criterion LI** to quantify the results shown in Fig 4. In this performance analysis, mean values of LI and their corresponding standard deviations were used to quantify the efficiency (depends on LI values) and consistency (depends on standard deviation values), respectively, of the scout functions and time-frequency decomposition algorithms. We conducted three analyses in TR case using different trial selections to evaluate the consistency of the methods based on the obtained results. Fig 5 depicts the mean values of the criterion LI on TR case for different scout functions (MEAN, MAX and PCA) and time-frequency decomposition algorithms (WT and HT). We performed the Pearson’s correlation analysis to describe the interplays between the variability of each factor and the different factors/combinations. We noted that the combinations with same scout function have lower correlation values (0.194 for Wavelet/PCA vs. Hilbert/PCA and Wavelet/MAX vs. Hilbert/MAX) than the combinations with same time-frequency decomposition algorithms (0.979 for Hilbert/PCA vs. Hilbert/MAX and 0.986 for Wavelet/PCA vs. Wavelet/MAX). The correlation values show that time-frequency decomposition algorithms have higher variability and therefore, the time-frequency decomposition algorithms strongly impact the LI values as compared to the scout functions.

**Fig 5.**
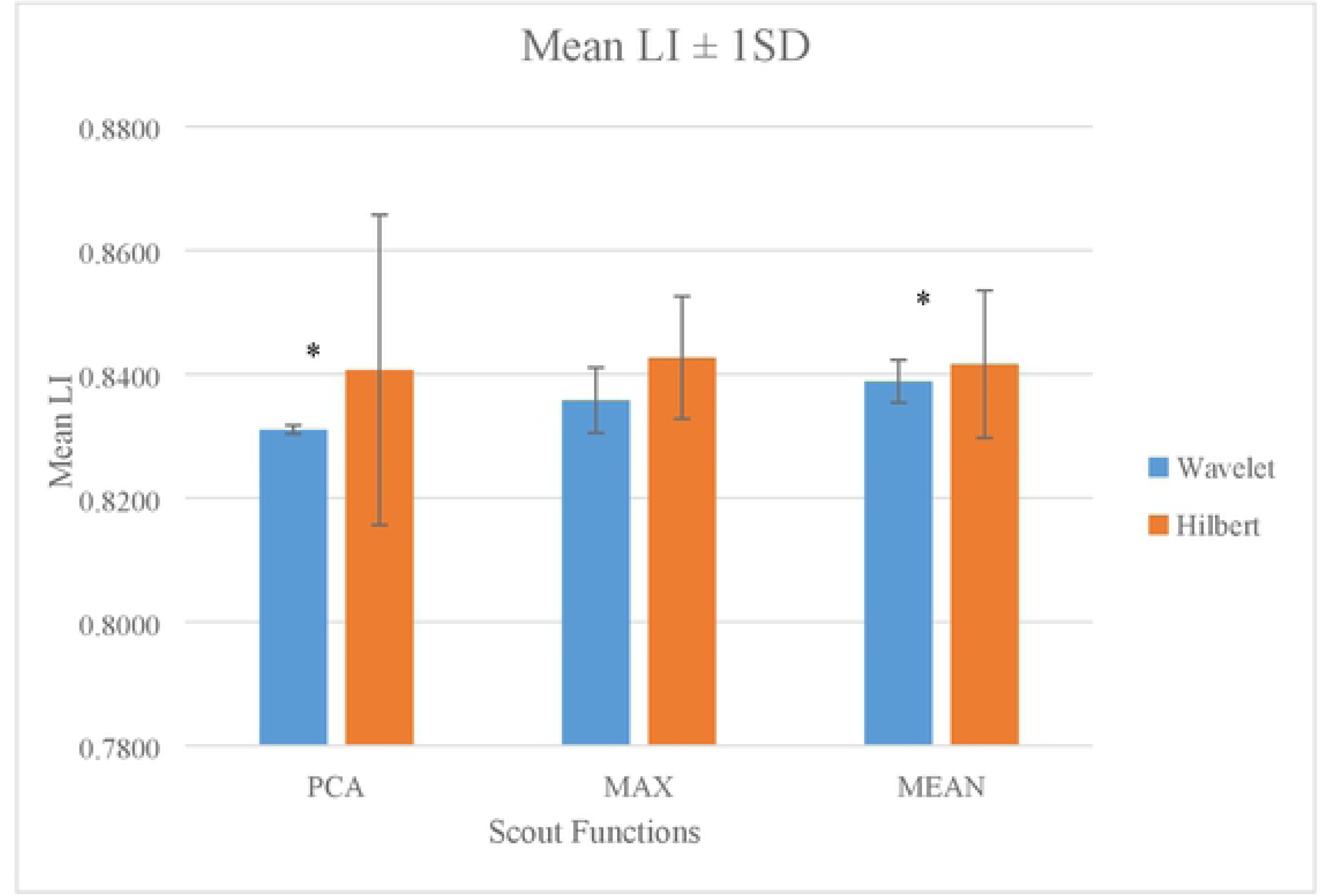
Comparison of averaged LI values between different time-frequency decomposition algorithms for different scout functions. Asterisk above the bars indicates significant difference (P≤S0.05). (LI: Localization Index; SD: Standard Deviation).

As observed, regardless of scout functions, HT yields slightly higher mean LI values than WT. Theoretically, HT yields slightly higher efficiency than WT based on the connectivity estimation over expected ROIs (based on literatures) using PLV. However, as we can observe, the difference between these two time-frequency decomposition algorithms in term of LI are minor (0.01 in PCA; 0.007 in MAX; 0.003 in MEAN). We performed Student’s t-test. The results show no significant difference between WT and HT in scout functions of PCA, MAX and MEAN. Application of both algorithms produces similar results with minor difference [69,90,91]. Therefore, we can conclude that application of both algorithms on connectivity estimation using PLV is efficient as the result shows that PLV is able to localize more than 80% of significant connections using both algorithms to extract the instantaneous phases of the regional time series. However, in terms of consistency, WT is better. Based on the results, we observe that WT is more consistent in extracting instantaneous phases for connectivity estimation as the standard deviation values are lower. As mentioned earlier, we conducted the analysis three times on same methodology by different trial selections in TR case. The standard deviation represents the fluctuations or more precisely, variability of the factors across different selected trials on the same case. In this situation, WT is more consistent than HT across the different trials. Therefore, we conclude that WT performed better in instantaneous phase extractions for connectivity estimation for visual oddball paradigm.

Hence time-frequency decomposition algorithms have stronger effect on mean LI values, whereas scout functions have minor impact on the mean LI values. Combinations of scout functions (PCA, MAX, and MEAN) with same time-frequency decomposition algorithm (HT) provide the higher mean LI value. The result indicates that these combinations have higher efficiency in connectivity estimation over expected ROIs using PLV. Among these combinations, MAX with HT provides highest averaged LI value (LI=0.843). However, some minor differences are observed among these combinations. According to the results of the Student’s t-test, these minor differences are insignificant. Hence, in term of efficiency, PCA, MAX and MEAN with HT provides similar results with minor differences. Nevertheless, in term of consistency, MAX/HT is better as the standard deviation is smaller among these three combinations. Therefore, the application of HT in conjunction with scout function MAX yields better outcome than the other two combinations.

On the other hand, combinations of WT with three scout functions produce lower mean LI values than combinations of HT with the three scout functions. In these three combinations, WT/MEAN gives higher mean LI value than others. However, the differences are still minor. Thus, we performed Student’s t-test. The results demonstrate that WT/PCA and WT/MEAN pairs are significantly different (p<0.05). WT/MEAN pair provides higher efficiency in connectivity estimation. However, in terms of consistency, WT/PCA pair is better.

Hence we can conclude that WT is better than HT in terms of consistency of the results [92]. WT with MEAN scout function gives higher efficiency in connectivity estimation. However, WT with PCA pair has better consistency than other combinations.

Now we discuss the outcomes obtained from the quantitative comparison of the performance of the time-frequency decomposition algorithms and scout functions on FC estimation, based on the **R criterion**. The R criterion is known as a ‘local’ indicator. The R values reflect the distribution of estimated links in each predefined ROI obtained from literature. R is known to be important as the brain regions activated during the visual oddball paradigm (consists of P300 and motor response) are supposed to be distinct. Moreover, the estimated network is known to be dependent on the time-frequency decomposition algorithms and scout functions.

In Fig 6, the straight line curves of the R values obtained for the two time-frequency decomposition algorithms are superimposed and plotted for each scout function. Results indicate that all signal processing algorithms identified a comparable percentage of significant connections for all scout functions. As shown in Fig 6 the SD values of WT/Scout functions is mostly lower than the HT/Scout functions. Only few SD values in WT/MAX and WT/MEAN pairs are slightly higher. (For examples: LF, LT and RT in WT/MAX pair; LF, LC, LT, RT and LP in WT/MEAN pair). However, the WT is still considered as a consistent algorithm to extract instantaneous phases for. This result is correlated with the result in the preceding part for LI calculation.

**Fig 6.**
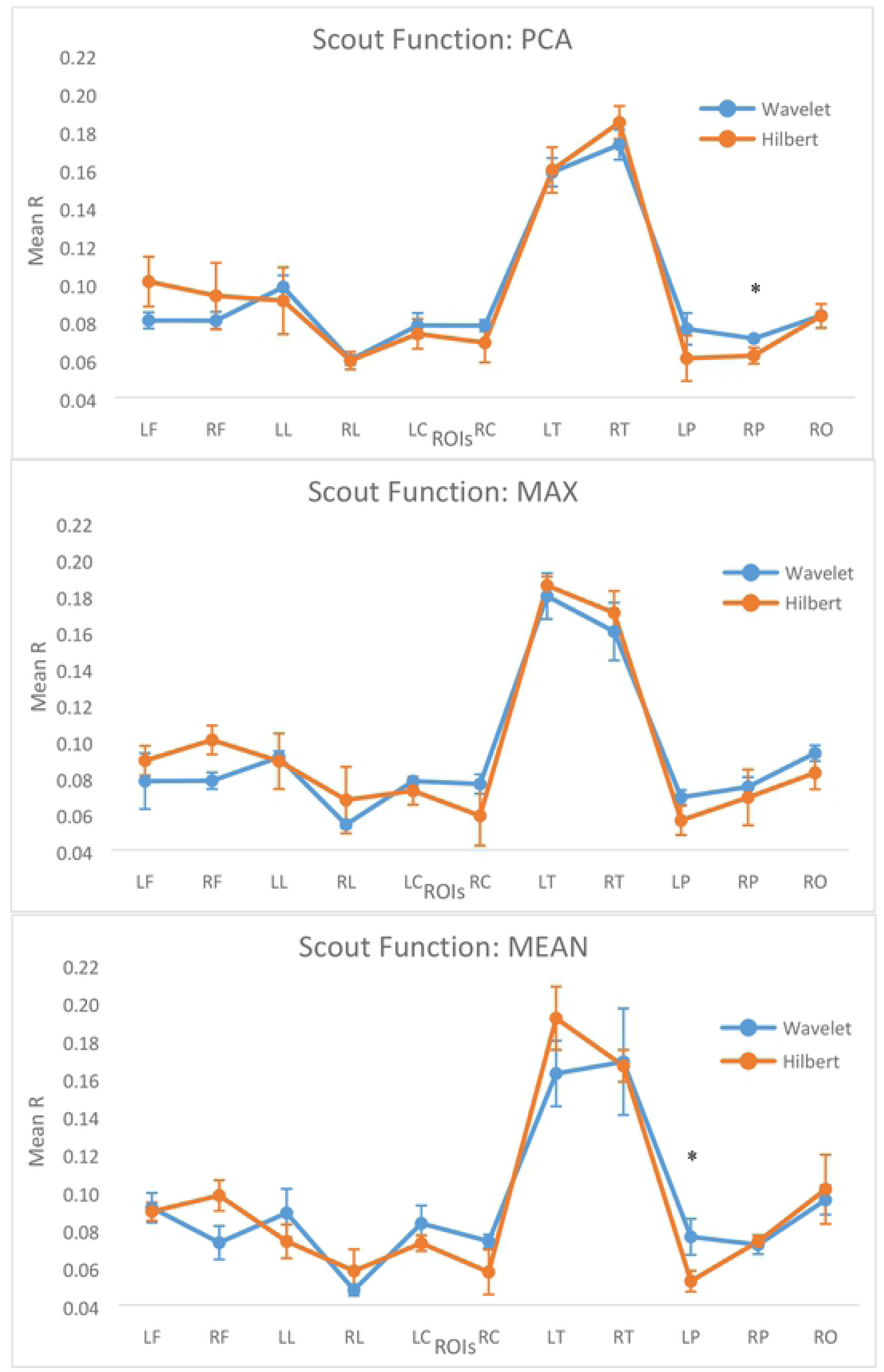
The mean and standard deviations of the R values (ratio of the identified connections within each ROI) obtained for the different scout functions.

Student’s t test was carried out to statistically compare the differences between different factors (WT/PCA vs. HT/PCA; WT/MAX vs. HT/MAX; WT/MEAN vs. HT/MEAN) in 4 different ROIs (LC; RC; LP; RP) to identify the significant pairs of factors with higher efficiency. These 4 ROIs are significant in oddball tasks [93-96]. Neural activity in LP and RP corresponds to P300 (decision making) while neural activity in LC and RC corresponds to motor planning and integration.

As shown in Fig 6, combinations of WT with 3 scout functions yield higher averaged R values as compared to combinations of HT with 3 scout functions. WT/PCA pair yields higher average R values on the 4 ROIs than HT/PCA pair. Results indicate no significant differences between the combinations in LC, RC and LP. However, a significant difference is observed in RP (p=0.023). On the other hand, as depicted in Fig 6, Wavelet/MAX gives higher average R values than Hilbert/MAX as well over the ROIs of LC, RC, LP and RP. The differences are still minor between these two combinations. Statistical outcomes demonstrate that no significant differences are observed on these two pairs of factors. Lastly, we applied Student’s t test on the combinations of WT/ MEAN and HT/MEAN. Results reveal that the differences are not between the combinations in LC, RC and RP. However, a significant difference is observed in LP (p=0.021).

Based on the statistical results, we note that WT/PCA pair yields greater R value than Hilbert/PCA over parietal region (RP) and the difference is significant. Moreover, WT/MEAN pair yields greater R value than Hilbert/MEAN over parietal region (LP) as well and the difference is significant. According to other researches, activity over parietal region is essentially elicited by P300 response which is a significant oddball event [97,98]. Thus, we believe that WT/PCA and WT/MEAN pairs provide good performance for PLV to localize the connectivity triggered by P300 response.

By combining the analysis from LI and R, we can say that WT is more consistent than HT. Moreover, we also realized that WT/PCA and WT/MEAN pairs have high performance. Hence the results of Wavelet/PCA pair is more consistent. Besides that, PLV also was able to localize more than 80% of networks (LI=0.831) using WT/PCA pair. The efficiency of this pair is acceptable and the consistency of this pair is better. Therefore, we conclude that WT/PCA is adequate and a good choice for visual Oddball task.

We performed the comparisons between TR and NTNR cases in order to validate the performance of the WT/PCA pair based on LI and R criterions. Moreover, we would like to analyse the differences of TR versus NTNR in terms of FC using R criterion.

Fig 7 shows the distribution of the estimated brain functional networks on cortical surface for TR and NTNR cases. As illustrated, the difference in terms of estimated connectivity among both cases is obvious. Consequently, target and non-target stimuli elicited different brain networks during oddball task. As depicted in Fig 7, the difference in the estimated networks is illustrated on the cortical surface. Furthermore, the quantitative analysis is required to evaluate the difference is done using LI and R parameters.

**Fig 7.**
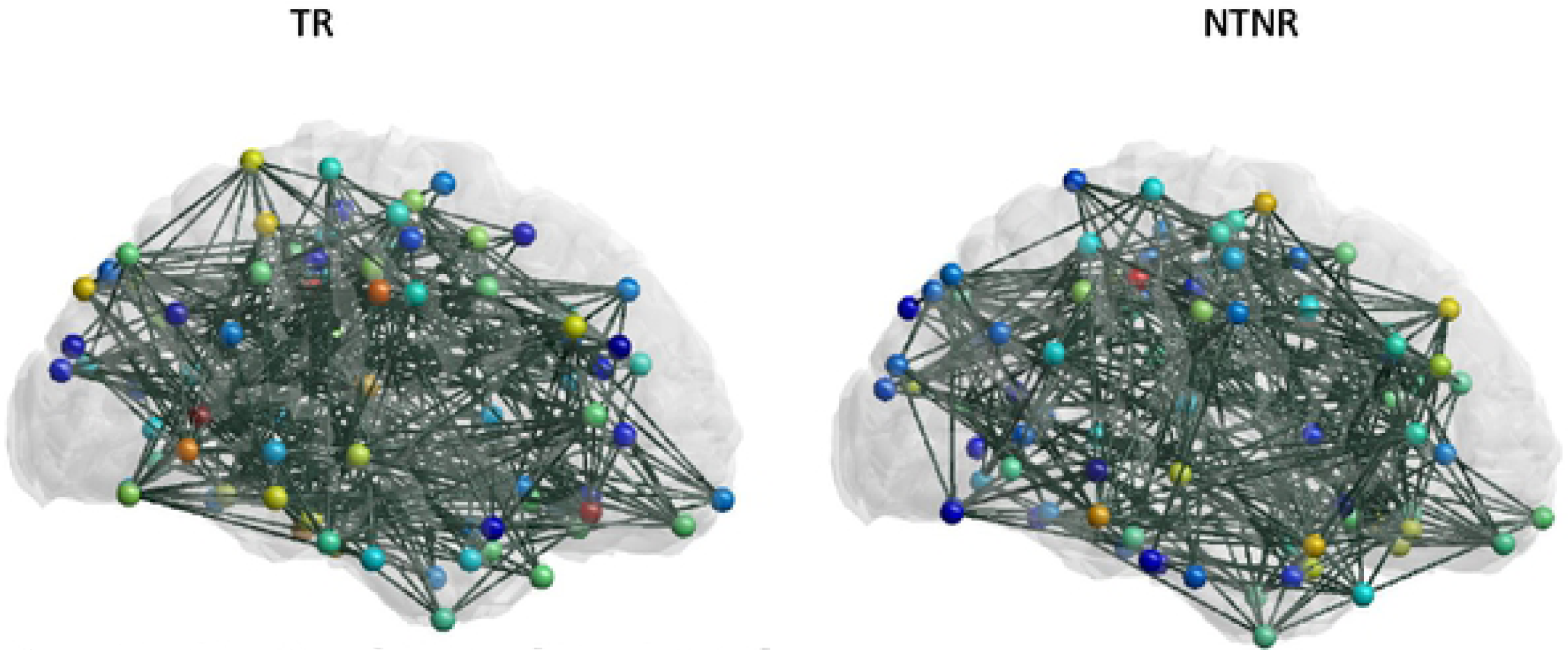
Cortical functional networks estimated using PLY based on the instantaneous phases extracted by Wavelet Transform/PCA pair for two oddball cases. Left panel: TR case; Right panel: NTNR case

Fig 8 shows the mean LI value of WT/PCA for TR and NTNR cases. It is observed that the PLV is able to localize denser networks over the predefined ROIs in TR case which are significant for visual oddball event including P300, motor response and others elicited by target stimuli. In our case, we observe that TR case has greater LI than NTNR case. In other words, target stimuli triggered higher brain connectivity. We performed statistical test (ANOVA) to evaluate the difference between these two cases. The TR case is significantly different (p=0.002) from NTNR case in terms of average LI values. Thus, we conclude that TR case triggered denser networks as compared to NTNR case [50,99]. Moreover, WT/PCA is sufficient for PLV to use as a connectivity estimation modality for visual oddball paradigm.

**Fig 8.**
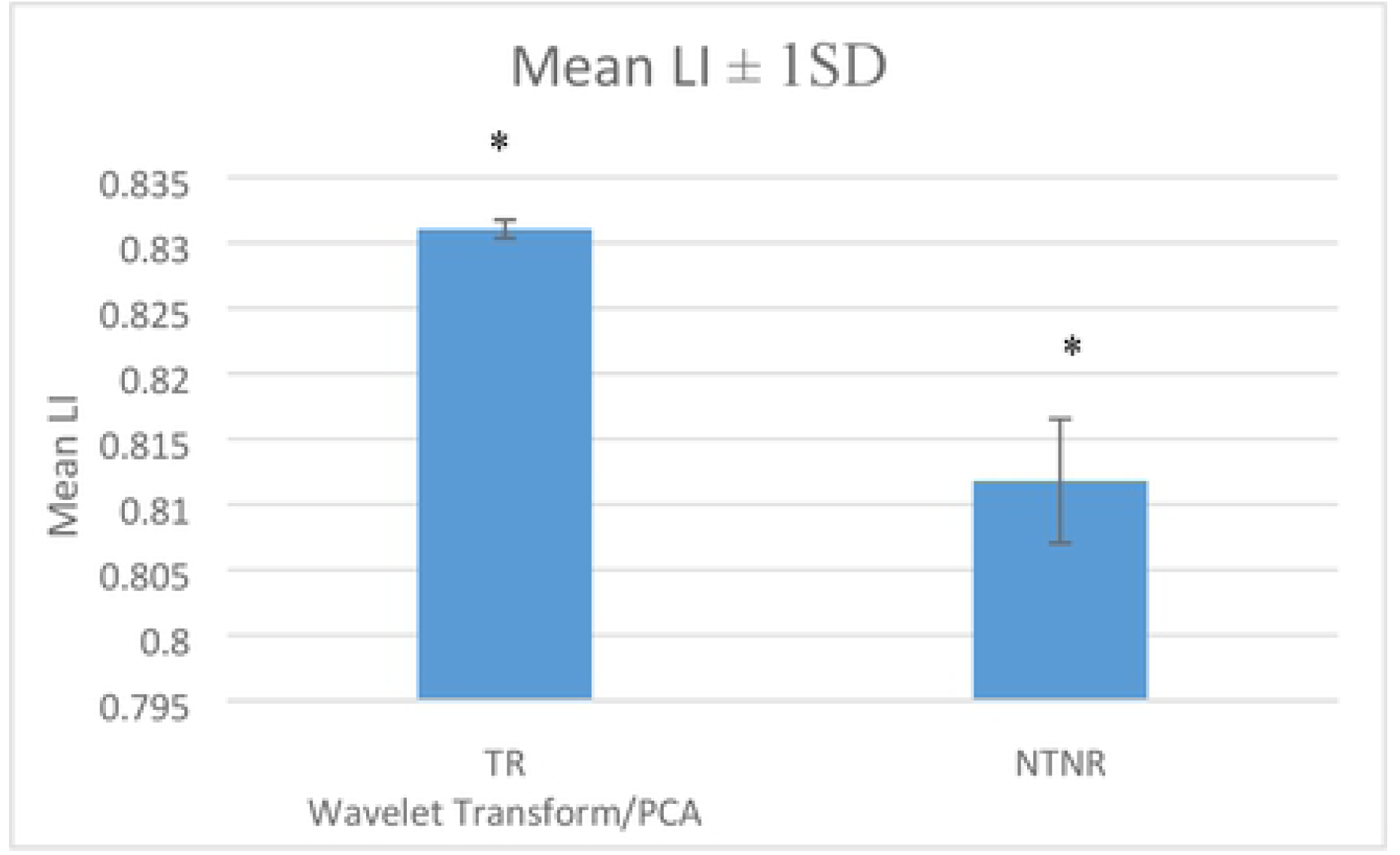
Comparison of mean LI between TR and NTNR cases. Asterisk above the bars indicates significant difference (P≤S0.05). LI: Localization Index; SD: Standard Deviation.

So far, we compared the difference between TR and NTNR case using ‘global’ indicator. Now, we will analyse the differences between TR and NTNR cases within ROIs using ‘local’ indicator, criterion R. The comparisons of R values between two cases were plotted on a line graph shown in Fig 9. As seen the major differences between TR and NTNR cases are observed within majority of ROIs especially LL, RL, LC, RC, LP, RP and LT. TR case has denser cortical connectivity within these ROIs than NTNR case. ANOVA tests were carried out for each ROI to statistically compare the difference between both cases within each ROI.

**Fig 9.**
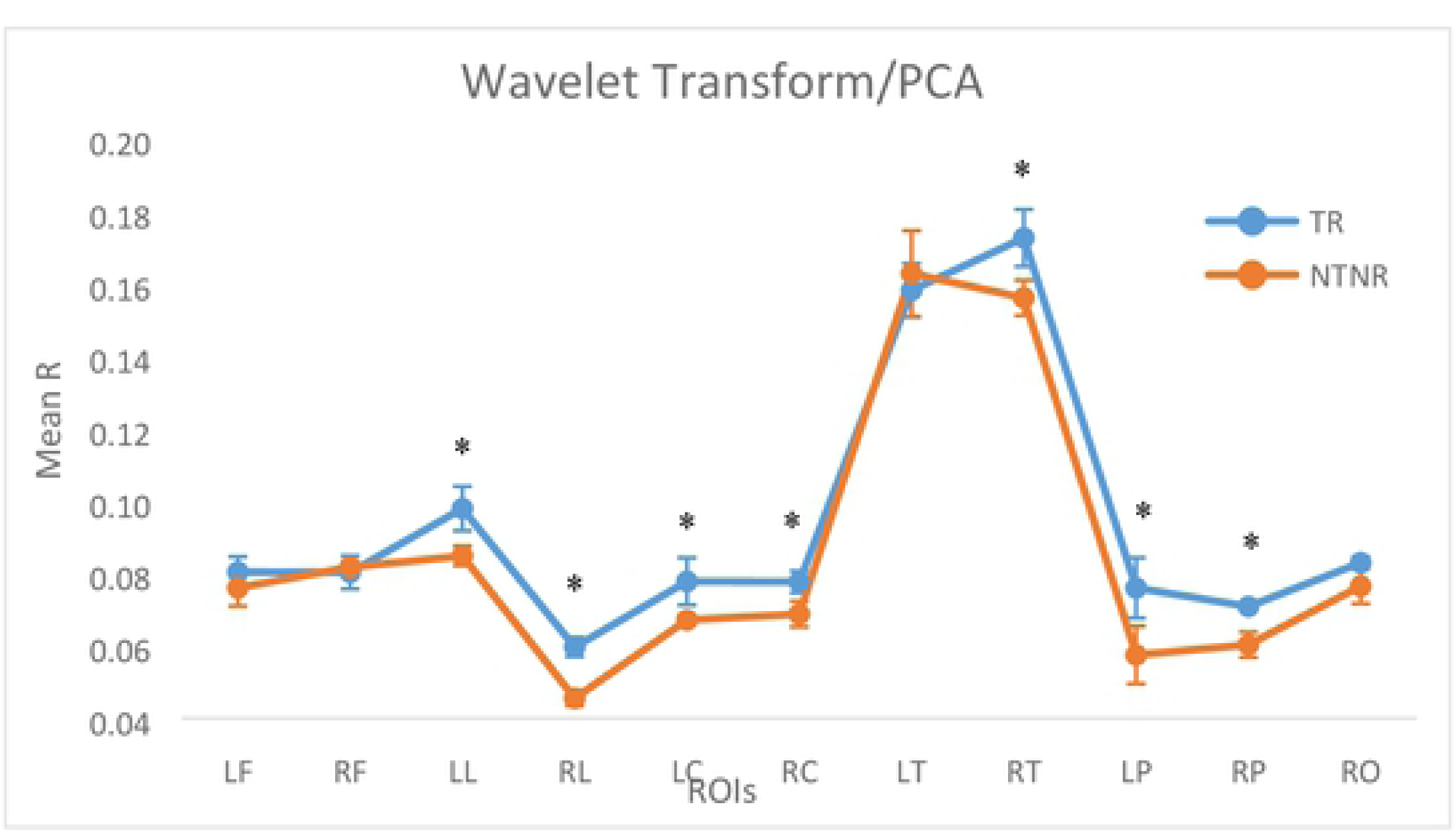
Comparison of mean R between TR and NTNR cases. Asterisk above the lines indicates significant difference (P<0.05). R: ratio of the identified connections for within each ROI; SD: Standard Deviation.

For ROIs in limbic (LL & RL) regions, target stimuli elicited greater cortical activity. Based on ANOVA the TR case is significantly different than NTNR case for R values within LL and RL (p=0.028 & p=0.002 respectively). According to Destrieux atlas, Cingulate cortices (Cingulate gyrus and sulcus) are included in the limbic regions. Our result show that target stimuli elicited denser networks over bilateral limbic regions than non-target stimuli which is in line with other researches [86,100,101]. These cortical activities are mostly related to P300 response (decision making for the response to target stimuli) [102-105] as well as target stimuli detection [106]. Therefore, TR case is different with NTNR case in terms of the estimated network over limbic regions as the target detection process and P300 response are present in TR case, but absent in NTNR and the difference is significant.

For ROIs in central (LC & RC) regions, the difference between TR and NTNR cases in terms of R is obvious. Greater central cortical activity is observed in TR case than in NTNR. Results of the statistical test shows that TR case is different from NTNR case in terms of the corresponding R values within these two ROIs and the differences are significant (LC: p=0.05; RC: p=0.03). According to our oddball paradigm, instead of non-target stimuli, the subjects provided motor response to the target stimuli. Thus, higher cortical activity is found over central regions in TR case than NTNR case. Our result is correlated with other studies [76,100,105,107-110]. Therefore, TR case is significantly different from NTNR in terms of cortical network over central regions.

For ROI in right temporal (RT) lobe, superior temporal activity is observed in TR case. ANOVA test indicates that the difference between TR and NTNR cases in terms of R values are significant (p=0.036) within this ROI. Some studies reported that cortical connectivity distributed over temporal regions corresponds to visual target detection [111] and P300 response [102,112,113]. Moreover, in [103,114], the authors reported that right temporal region is more dominant than left temporal during P300 response. Hence, TR case is different from NTNR case in terms of the connectivity over right temporal region for P300 response.

Lastly, in parietal (LP & RP) regions, higher parietal activity is noticed in TR case. The results of statistical tests denote that both cases are significantly distinct with each other in LP (p=0.049) and RP (p=0.007) in terms of their designated R values. Denser bilateral cortical activity is observed over parietal regions in TR case that corresponds to P300 response [105,115-118] and motor coordination for button pressing [76,119,120]. Thus, NTNR case has lower bilateral parietal activity as no P300 and motor responses are required for non-target stimuli.

## Conclusion

In this study our aim is to observe the FC for visual oddball task using the source space. We used wMNE method to reconstruct the sources for data acquired by EEG for visual oddball task. We applied three different scout functions (MEAN, MAX and PCA) to generate the regional time series signals. We applied two time-frequency decomposition algorithms (HT and WT) to represent the regional time series signals into complex functions. We extracted instantaneous phases from the complex form of regional time series signals and estimated the FC using PLV. The connectivity graphs were proportionally thresholded to retain 10% of strongest networks. We evaluated the performance of the scout functions/time-frequency decomposition algorithms pairs based on LI and R. Lastly, we compared the differences between TR and NTNR cases based on the LI and R.

Our results demonstrate that time-frequency decomposition algorithms have higher impact on the LI values than the scout functions. In addition, WT is better than HT in terms of the consistency of LI. All pairs show good efficiency in connectivity estimation as all pairs yield more than 80% of LI. However, WT/PCA pair is more consistent than others. Moreover, WT/PCA is capable to estimate the connectivity within parietal region which corresponds to P300 response. Lastly, we observe the differences in connectivity between TR and NTNR cases over parietal, central, right temporal and limbic regions which correspond to target detection, P300 response and motor response.

In conclusion, the outcome of the connectivity estimation might be affected by scout functions/time-frequency algorithm pairs. Consequently, WT/PCA is the best choice for visual oddball task as the scout function to generate regional time series signals and time-frequency decomposition algorithms to transform the signals into gamma band for instantaneous phase extraction. The TR and NTNR cases are different in terms of FC. Greater R values are observed over the regions which correspond to P300 and motor response.

The performance of the combinations of scout functions/time-frequency decomposition algorithms have not been evaluated so far in the literature. We found that the WT/PCA is best for visual oddball task. We believe that PCA is superior for generation of regional time series signals and WT is superior for time-frequency decomposition for extraction of instantaneous phases. On the other hand, we found that higher connectivity over parietal, temporal, central region and limbic regions are significant for P300 response. Thus, we believe that this discovery of P300 response could be used as an electrophysiological marker to distinguish the healthy individuals and the subjects with mild cognitive impairment diseases as well as the marker for the diagnosis and prediction of mild cognitive impairment disorders [96,121-123].

## References

1. Beaty RE, Chen Q, Christensen AP, Qiu J, Silvia PJ, Schacter DL. Brain networks of the imaginative mind: Dynamic functional connectivity of default and cognitive control networks relates to openness to experience. Hum Brain Mapp. 2018;39(2):811–21.

2. Giraldo-Chica M, Rogers BP, Damon SM, Landman BA, Woodward ND. Prefrontal-thalamic anatomical connectivity and executive cognitive function in schizophrenia. Biol Psychiatry. 2018;83(6):509–17.

3. Cléry H, Andersson F, Fonlupt P, Gomot M. Brain correlates of automatic visual change detection. Neuroimage. 2013;75:117–22.

4. Fornito A, Yoon J, Zalesky A, Bullmore ET, Carter CS. General and specific functional connectivity disturbances in first-episode schizophrenia during cognitive control performance. Biol Psychiatry. 2011;70(1):64–72.

5. Lynall M-E, Bassett DS, Kerwin R, McKenna PJ, Kitzbichler M, Muller U, et al. Functional connectivity and brain networks in schizophrenia. J Neurosci. 2010;30(28):9477–87.

6. Van Den Heuvel MP, Pol HEH. Exploring the brain network: a review on resting-state fMRI functional connectivity. Eur Neuropsychopharmacol. 2010;20(8):519–34.

7. Englander ZA, Sun J, Case L, Mikati MA, Kurtzberg J, Song AW. Brain structural connectivity increases concurrent with functional improvement: evidence from diffusion tensor MRI in children with cerebral palsy during therapy. NeuroImage Clin. 2015;7:315–24.

8. Sitek KR, Cai S, Beal DS, Perkell JS, Guenther FH, Ghosh SS. Decreased cerebellar-orbitofrontal connectivity correlates with stuttering severity: whole-brain functional and structural connectivity associations with persistent developmental stuttering. Front Hum Neurosci. 2016;10:190.

9. Ewert S, Plettig P, Li N, Chakravarty MM, Collins DL, Herrington TM, et al. Toward defining deep brain stimulation targets in MNI space: a subcortical atlas based on multimodal MRI, histology and structural connectivity. Neuroimage. 2018;170:271–82.

10. Mueller F, Musso F, London M, de Boer P, Zacharias N, Winterer G. Pharmacological fMRI: Effects of subanesthetic ketamine on resting-state functional connectivity in the default mode network, salience network, dorsal attention network and executive control network. NeuroImage Clin. 2018;19:745–57.

11. Hindriks R, Adhikari MH, Murayama Y, Ganzetti M, Mantini D, Logothetis NK, et al. Can sliding-window correlations reveal dynamic functional connectivity in resting-state fMRI? Neuroimage. 2016;127:242–56.

12. Zhang S, Chiang-shan RL. Functional connectivity mapping of the human precuneus by resting state fMRI. Neuroimage. 2012;59(4):3548–62.

13. Tagliazucchi E, Laufs H. Multimodal imaging of dynamic functional connectivity. Front Neurol. 2015;6:10.

14. Vitali P, Di Perri C, Vaudano AE, Meletti S, Villani F. Integration of multimodal neuroimaging methods: a rationale for clinical applications of simultaneous EEG-fMRI. Funct Neurol. 2015;30(1):9.

15. Coito A, Michel CM, van Mierlo P, Vulliémoz S, Plomp G. Directed functional brain connectivity based on EEG source imaging: methodology and application to temporal lobe epilepsy. IEEE Trans Biomed Eng. 2016;63(12):2619–28.

16. Mheich A, Hassan M, Dufor O, Khalil M, Berrou C, Wendling F. Spatiotemporal analysis of brain functional connectivity. In: 6th European Conference of the International Federation for Medical and Biological Engineering. 2015. p. 934–7.

17. Sakkalis V. Review of advanced techniques for the estimation of brain connectivity measured with EEG/MEG. Comput Biol Med. 2011;41(12):1110–7.

18. David O, Cosmelli D, Hasboun D, Garnero L. A multitrial analysis for revealing significant corticocortical networks in magnetoencephalography and electroencephalography. Neuroimage. 2003;20(1):186–201.

19. Somon B, Campagne A, Delorme A, Berberian B. Evaluation of performance monitoring ERPs through difficulty manipulation in a response-feedback paradigm. Brain Res. 2019;1704:196–206.

20. Burle B, Spieser L, Roger C, Casini L, Hasbroucq T, Vidal F. Spatial and temporal resolutions of EEG: Is it really black and white? A scalp current density view. Int J Psychophysiol. 2015;97(3):210–20.

21. Hashiguchi K, Morioka T, Yoshida F, Miyagi Y, Nagata S, Sakata A, et al. Correlation between scalprecorded electroencephalographic and electrocorticographic activities during ictal period. Seizure. 2007;16(3):238–47.

22. Stam CJ van, Van Straaten ECW. The organization of physiological brain networks. Clin Neurophysiol. 2012;123(6):1067–87.

23. Lachaux J-P, Rodriguez E, Martinerie J, Varela FJ, others. Measuring phase synchrony in brain signals. Hum Brain Mapp. 1999;8(4):194–208.

24. Nolte G, Bai O, Wheaton L, Mari Z, Vorbach S, Hallett M. Identifying true brain interaction from EEG data using the imaginary part of coherency. Clin Neurophysiol. 2004;115(10):2292–307.

25. Bola Michałand Sabel BA. Dynamic reorganization of brain functional networks during cognition. Neuroimage. 2015;114:398–413.

26. Hassan M, Dufor O, Merlet I, Berrou C, Wendling F. EEG source connectivity analysis: from dense array recordings to brain networks. PLoS One. 2014;9(8):e105041.

27. Schoffelen J-M, Gross J. Source connectivity analysis with MEG and EEG. Hum Brain Mapp. 2009;30(6):1857–65.

28. Bradley A, Yao J, Dewald J, Richter C-P. Evaluation of electroencephalography source localization algorithms with multiple cortical sources. PLoS One. 2016;11(1):e0147266.

29. Becker H, Albera L, Comon P, Haardt M, Birot G, Wendling F, et al. EEG extended source localization: tensor-based vs. conventional methods. Neuroimage. 2014;96:143–57.

30. Jatoi MA, Kamel N, Malik AS, Faye I, Begum T. A survey of methods used for source localization using EEG signals. Biomed Signal Process Control. 2014;11:42–52.

31. Kaiboriboon K, Lüders HO, Hamaneh M, Turnbull J, Lhatoo SD. EEG source imaging in epilepsy— practicalities and pitfalls. Nat Rev Neurol. 2012;8(9):498.

32. Bastos AM, Schoffelen J-M. A tutorial review of functional connectivity analysis methods and their interpretational pitfalls. Front Syst Neurosci. 2016;9:175.

33. Greenblatt RE, Pflieger ME, Ossadtchi AE. Connectivity measures applied to human brain electrophysiological data. J Neurosci Methods. 2012;207(1):1–16.

34. Li F, Yi C, Jiang Y, Liao Y, Si Y, Yao D, et al. The Construction of Large-Scale Cortical Networks for P300 From Scalp EEG. IEEE Access. 2018;6:68498–506.

35. Liu Q, Ganzetti M, Wenderoth N, Mantini D. Detecting large-scale brain networks using EEG: impact of electrode density, head modeling and source localization. Front Neuroinform. 2018;12:4.

36. Supp GG, Schlögl A, Trujillo-Barreto N, Müller MM, Gruber T. Directed cortical information flow during human object recognition: analyzing induced EEG gamma-band responses in brain’s source space. PLoS One. 2007;2(8):e684.

37. Astolfi L, Cincotti F, Mattia D, Babiloni C, Carducci F, Basilisco A, et al. Assessing cortical functional connectivity by linear inverse estimation and directed transfer function: simulations and application to real data. Clin Neurophysiol. 2005;116(4):920–32.

38. Hämäläinen MS, Ilmoniemi RJ. Interpreting magnetic fields of the brain: minimum norm estimates. Med Biol Eng Comput. 1994;32(1):35–42.

39. Lin F-H, Witzel T, Ahlfors SP, Stufflebeam SM, Belliveau JW, Hämäläinen MS. Assessing and improving the spatial accuracy in MEG source localization by depth-weighted minimum-norm estimates. Neuroimage. 2006;31(1):160–71.

40. Pascual-Marqui RD, Michel CM, Lehmann D. Low resolution electromagnetic tomography: a new method for localizing electrical activity in the brain. Int J Psychophysiol. 1994;18(1):49–65.

41. Pascual-Marqui RD, others. Standardized low-resolution brain electromagnetic tomography (sLORETA): technical details. Methods Find Exp Clin Pharmacol. 2002;24(Suppl D):5–12.

42. López ME, Pusil S, Pereda E, Maestú F, Barceló F. Dynamic low frequency EEG phase synchronization patterns during proactive control of task switching. Neuroimage. 2019;186:70–82.

43. Kiat JE. Assessing cross-modal target transition effects with a visual-auditory oddball. Int J Psychophysiol. 2018;129:58–66.

44. Molina V, Bachiller A, de Luis R, Lubeiro A, Poza J, Hornero R, et al. Topography of activation deficits in schizophrenia during P300 task related to cognition and structural connectivity. Eur Arch Psychiatry Clin Neurosci. 2018;1–10.

45. Shim M, Kim D-W, Lee S-H, Im C-H. Disruptions in small-world cortical functional connectivity network during an auditory oddball paradigm task in patients with schizophrenia. Schizophr Res. 2014;156(2–3):197–203.

46. Brazier MAB, Casby JU. Crosscorrelation and autocorrelation studies of electroencephalographic potentials. Electroencephalogr Clin Neurophysiol. 1952;4(2):201–11.

47. Brazier MAB. Studies of the EEG activity of limbic structures in man. Electroencephalogr Clin Neurophysiol. 1968;25(4):309–18.

48. Mars NJ, da Silva Lopes FH. Propagation of seizure activity in kindled dogs. Electroencephalogr Clin Neurophysiol. 1983;56(2):194–209.

49. Rosenblum M, Pikovsky A, Kurths J, Schäfer C, Tass PA. Phase synchronization: from theory to data analysis. In: Handbook of biological physics. Elsevier; 2001. p. 279–321.

50. Thee KW, Nisar H, Soh CS. Graph Theoretical Analysis of Functional Brain Networks in Healthy Subjects: Visual Oddball Paradigm. IEEE Access. 2018;6:64708–27.

51. Lim SH, Nisar H, Yap VV, Shim S-O. Tracking of electroencephalography signals across brain lobes using motion estimation and cross-correlation. J Electron Imaging. 2015;24(6):61106.

52. Mheich A, Hassan M, Khalil M, Berrou C, Wendling F. A new algorithm for spatiotemporal analysis of brain functional connectivity. J Neurosci Methods. 2015;242:77–81.

53. Wang SH, Lobier M, Siebenhühner F, Puoliväli T, Palva S, Palva JM. Hyperedge bundling: A practical solution to spurious interactions in MEG/EEG source connectivity analyses. Neuroimage. 2018;173:610–22.

54. Cohen MX, Ridderinkhof KR. EEG source reconstruction reveals frontal-parietal dynamics of spatial conflict processing. PLoS One. 2013;8(2):e57293.

55. Hillebrand A, Barnes GR, Bosboom JL, Berendse HW, Stam CJ. Frequency-dependent functional connectivity within resting-state networks: an atlas-based MEG beamformer solution. Neuroimage. 2012;59(4):3909–21.

56. Lehmann D, Faber PL, Tei S, Pascual-Marqui RD, Milz P, Kochi K. Reduced functional connectivity between cortical sources in five meditation traditions detected with lagged coherence using EEG tomography. Neuroimage. 2012;60(2):1574–86.

57. Hassan M, Merlet I, Mheich A, Kabbara A, Biraben A, Nica A, et al. Identification of interictal epileptic networks from dense-EEG. Brain Topogr. 2017;30(1):60–76.

58. Lai M, Demuru M, Hillebrand A, Fraschini M. A comparison between scalp-and source-reconstructed EEG networks. Sci Rep. 2018;8(1):12269.

59. Kabbara A, Falou WEL, Khalil M, Wendling F, Hassan M. The dynamic functional core network of the human brain at rest. Sci Rep. 2017;7(1):2936.

60. Hassan M, Benquet P, Biraben A, Berrou C, Dufor O, Wendling F. Dynamic reorganization of functional brain networks during picture naming. Cortex. 2015;73:276–88.

61. Aviyente S, Tootell A, Bernat EM. Time-frequency phase-synchrony approaches with ERPs. Int J Psychophysiol. 2017;111:88–97.

62. Destrieux C, Fischl B, Dale A, Halgren E. Automatic parcellation of human cortical gyri and sulci using standard anatomical nomenclature. Neuroimage. 2010;53(1):1–15.

63. Tadel F, Baillet S, Mosher JC, Pantazis D, Leahy RM. Brainstorm: a user-friendly application for MEG/EEG analysis. Comput Intell Neurosci. 2011;2011:8.

64. Lim SH, Nisar H, Thee KW, Yap VV. A novel method for tracking and analysis of EEG activation across brain lobes. Biomed Signal Process Control. 2018;40:488–504.

65. Fonov V, Evans AC, Botteron K, Almli CR, McKinstry RC, Collins DL, et al. Unbiased average ageappropriate atlases for pediatric studies. Neuroimage. 2011;54(1):313–27.

66. Mazziotta JC, Toga AW, Evans A, Fox P, Lancaster J, others. A probabilistic atlas of the human brain: theory and rationale for its development. Neuroimage. 1995;2(2):89–101.

67. Mosher JC, Leahy RM, Lewis PS. EEG and MEG: forward solutions for inverse methods. IEEE Trans Biomed Eng. 1999;46(3):245–59.

68. Gramfort A, Papadopoulo T, Olivi E, Clerc M. OpenMEEG: opensource software for quasistatic bioelectromagnetics. Biomed Eng Online. 2010;9(1):45.

69. Le Van Quyen M, Foucher J, Lachaux J-P, Rodriguez E, Lutz A, Martinerie J, et al. Comparison of Hilbert transform and wavelet methods for the analysis of neuronal synchrony. J Neurosci Methods. 2001;111(2):83–98.

70. Qassim YT, Cutmore TRH, James DA, Rowlands DD. Wavelet coherence of EEG signals for a visual oddball task. Comput Biol Med. 2013;43(1):23–31.

71. Thee KW, Nisar H. Comparison of Brain Functional Networks for Subjects with Different Performance. In: 2018 IEEE Region 10 Humanitarian Technology Conference (R10-HTC). 2018. p. 1–6.

72. Sporns O. Structure and function of complex brain networks. Dialogues Clin Neurosci. 2013;15(3):247.

73. Fajkus J, Mikl M, Shaw DJ, Brázdil M. An fMRI investigation into the effect of preceding stimuli during visual oddball tasks. J Neurosci Methods. 2015;251:56–61.

74. Kim YY, Jung YS. Reduced frontal activity during response inhibition in individuals with psychopathic traits: An sLORETA study. Biol Psychol. 2014;97:49–59.

75. Machado S, Arias-Carrión O, Sampaio I, Bittencourt J, Velasques B, Teixeira S, et al. Source imaging of P300 visual evoked potentials and cognitive functions in healthy subjects. Clin EEG Neurosci. 2014;45(4):262–8.

76. Walz JM, Goldman RI, Carapezza M, Muraskin J, Brown TR, Sajda P. Simultaneous EEG--fMRI reveals a temporal cascade of task-related and default-mode activations during a simple target detection task. Neuroimage. 2014;102:229–39.

77. Akimoto Y, Kanno A, Kambara T, Nozawa T, Sugiura M, Okumura E, et al. Spatiotemporal dynamics of high-gamma activities during a 3-stimulus visual oddball task. PLoS One. 2013;8(3):e59969.

78. Warbrick T, Reske M, Shah NJ. Do EEG paradigms work in fMRI? Varying task demands in the visual oddball paradigm: Implications for task design and results interpretation. Neuroimage. 2013;77:177–85.

79. Warbrick T, Mobascher A, Brinkmeyer J, Musso F, Stoecker T, Shah NJ, et al. Nicotine effects on brain function during a visual oddball task: a comparison between conventional and EEG-informed fMRI analysis. J Cogn Neurosci. 2012;24(8):1682–94.

80. Bocquillon P, Bourriez J-L, Palmero-Soler E, Betrouni N, Houdayer E, Derambure P, et al. Use of swLORETA to localize the cortical sources of target-and distracter-elicited P300 components. Clin Neurophysiol. 2011;122(10):1991–2002.

81. Rubia K, Hyde Z, Halari R, Giampietro V, Smith A. Effects of age and sex on developmental neural networks of visual--spatial attention allocation. Neuroimage. 2010;51(2):817–27.

82. Brázdil M, Mikl M, Mareček R, Krupa P, Rektor I. Effective connectivity in target stimulus processing: a dynamic causal modeling study of visual oddball task. Neuroimage. 2007;35(2):827–35.

83. Babiloni F, Cincotti F, Babiloni C, Carducci F, Mattia D, Astolfi L, et al. Estimation of the cortical functional connectivity with the multimodal integration of high-resolution EEG and fMRI data by directed transfer function. Neuroimage. 2005;24(1):118–31.

84. Ardekani BA, Choi SJ, Hossein-Zadeh G-A, Porjesz B, Tanabe JL, Lim KO, et al. Functional magnetic resonance imaging of brain activity in the visual oddball task. Cogn Brain Res. 2002;14(3):347–56.

85. Kiehl KA, Laurens KR, Duty TL, Forster BB, Liddle PF. An event-related fMRI study of visual and auditory oddball tasks. J Psychophysiol. 2001;15(4):221.

86. Stevens AA, Skudlarski P, Gatenby JC, Gore JC. Event-related fMRI of auditory and visual oddball tasks. Magn Reson Imaging. 2000;18(5):495–502.

87. Clark VP, Fannon S, Lai S, Benson R, Bauer L. Responses to rare visual target and distractor stimuli using event-related fMRI. J Neurophysiol. 2000;83(5):3133–9.

88. Downar J, Crawley AP, Mikulis DJ, Davis KD. A multimodal cortical network for the detection of changes in the sensory environment. Nat Neurosci. 2000;3(3):277.

89. Kirino E, Belger A, Goldman-Rakic P, McCarthy G. Prefrontal activation evoked by infrequent target and novel stimuli in a visual target detection task: an event-related functional magnetic resonance imaging study. J Neurosci. 2000;20(17):6612–8.

90. Yan B, Miyamoto A. A comparative study of modal parameter identification based on wavelet and Hilbert--Huang transforms. Comput Civ Infrastruct Eng. 2006;21(1):9–23.

91. Bruns A. Fourier-, Hilbert-and wavelet-based signal analysis: are they really different approaches? J Neurosci Methods. 2004;137(2):321–32.

92. Li D, Li X, Cui D, Li Z. Phase synchronization with harmonic wavelet transform with application to neuronal populations. Neurocomputing. 2011;74(17):3389–403.

93. Van Dinteren R, Huster RJ, Jongsma MLA, Kessels RPC, Arns M. Differences in cortical sources of the event-related P3 potential between young and old participants indicate frontal compensation. Brain Topogr. 2018;31(1):35–46.

94. Zhou L, Wang G, Nan C, Wang H, Liu Z, Bai H. Abnormalities in P300 components in depression: an ERP-sLORETA study. Nord J Psychiatry. 2018;1–8.

95. Hong X, Wang Y, Sun J, Li C, Tong S. Segregating Top-Down Selective Attention from Response Inhibition in a Spatial Cueing Go/NoGo Task: An ERP and Source Localization Study. Sci Rep. 2017;7(1):9662.

96. Yang P, Fan C, Wang M, Fogelson N, Li L. The effects of changes in object location on object identity detection: A simultaneous EEG-fMRI study. Neuroimage. 2017;157:351–63.

97. Overbye K, Huster RJ, Walhovd KB, Fjell AM, Tamnes CK. Development of the P300 from childhood to adulthood: a multimodal EEG and MRI study. Brain Struct Funct. 2018;223(9):4337–49.

98. Naeije G, Vaulet T, Wens V, Marty B, Goldman S, De Tiège X. Multilevel cortical processing of somatosensory novelty: a magnetoencephalography study. Front Hum Neurosci. 2016;10:259.

99. Kabbara A, Khalil M, El-Falou W, Eid H, Hassan M. Functional Brain Connectivity as a New Feature for P300 Speller. PLoS One. 2016;11(1):e0146282.

100. Warbrick T, Mobascher A, Brinkmeyer J, Musso F, Richter N, Stoecker T, et al. Single-trial P3 amplitude and latency informed event-related fMRI models yield different BOLD response patterns to a target detection task. Neuroimage. 2009;47(4):1532–44.

101. Laurens KR, Kiehl KA, Liddle PF. A supramodal limbic-paralimbic-neocortical network supports goaldirected stimulus processing. Hum Brain Mapp. 2005;24(1):35–49.

102. Wynn JK, Jimenez AM, Roach BJ, Korb A, Lee J, Horan WP, et al. Impaired target detection in schizophrenia and the ventral attentional network: findings from a joint event-related potential--functional MRI analysis. NeuroImage Clin. 2015;9:95–102.

103. Campanella S, Bourguignon M, Peigneux P, Metens T, Nouali M, Goldman S, et al. BOLD response to deviant face detection informed by P300 event-related potential parameters: A simultaneous ERP--fMRI study. Neuroimage. 2013;71:92–103.

104. Shenhav A, Botvinick MM, Cohen JD. The expected value of control: an integrative theory of anterior cingulate cortex function. Neuron. 2013;79(2):217–40.

105. Li Y, Wang L-Q, Hu Y. Localizing P300 generators in high-density event-related potential with fMRI. Med Sci Monit. 2009;15(3):MT47-MT53.

106. Harsay HA, Spaan M, Wijnen JG, Ridderinkhof KR. Error awareness and salience processing in the oddball task: shared neural mechanisms. Front Hum Neurosci. 2012;6:246.

107. Ragazzoni A, Di Russo F, Fabbri S, Pesaresi I, Di Rollo A, Perri RL, et al. “Hit the missing stimulus”. A simultaneous EEG-fMRI study to localize the generators of endogenous ERPs in an omitted target paradigm. Sci Rep. 2019;9(1):3684.

108. Cieslik EC, Zilles K, Grefkes C, Eickhoff SB. Dynamic interactions in the fronto-parietal network during a manual stimulus--response compatibility task. Neuroimage. 2011;58(3):860–9.

109. Bénar C-G, Schön D, Grimault S, Nazarian B, Burle B, Roth M, et al. Single-trial analysis of oddball event-related potentials in simultaneous EEG-fMRI. Hum Brain Mapp. 2007;28(7):602–13.

110. Bledowski C, Prvulovic D, Goebel R, Zanella FE, Linden DEJ. Attentional systems in target and distractor processing: a combined ERP and fMRI study. Neuroimage. 2004;22(2):530–40.

111. Staffen W, Ladurner G, Höller Y, Bergmann J, Aichhorn M, Golaszewski S, et al. Brain activation disturbance for target detection in patients with mild cognitive impairment: an fMRI study. Neurobiol Aging. 2012;33(5):1002-e1.

112. Cui H, Xie X, Xu S, Yan H, Feng L, Hu Y. Source analysis of bimodal event-related potentials with auditory-visual stimuli. In: 2013 6th International IEEE/EMBS Conference on Neural Engineering (NER). 2013. p. 85–8.

113. Helenius P, Laasonen M, Hokkanen L, Paetau R, Niemivirta M. Neural correlates of late positivities associated with infrequent visual events and response errors. Neuroimage. 2010;53(2):619–28.

114. Bledowski C, Prvulovic D, Hoechstetter K, Scherg M, Wibral M, Goebel R, et al. Localizing P300 generators in visual target and distractor processing: a combined event-related potential and functional magnetic resonance imaging study. J Neurosci. 2004;24(42):9353–60.

115. Li F, Chen B, Li H, Zhang T, Wang F, Jiang Y, et al. The time-varying networks in P300: a task-evoked EEG study. IEEE Trans Neural Syst Rehabil Eng. 2016;24(7):725–33.

116. Li F, Liu T, Wang F, Li H, Gong D, Zhang R, et al. Relationships between the resting-state network and the P3: Evidence from a scalp EEG study. Sci Rep. 2015;5:15129.

117. Rusiniak M, Lewandowska M, Wolak T, Pluta A, Milner Rafałand Ganc M, Włodarczyk A, et al. A modified oddball paradigm for investigation of neural correlates of attention: a simultaneous ERP--fMRI study. Magn Reson Mater Physics, Biol Med. 2013;26(6):511–26.

118. Babiloni C, Del Percio C, Triggiani AI, Marzano N, Valenzano A, De Rosas M, et al. Frontal-parietal responses to “oddball” stimuli depicting “fattened” faces are increased in successful dieters: An electroencephalographic study. Int J Psychophysiol. 2011;82(2):153–66.

119. Taylor KS, Seminowicz DA, Davis KD. Two systems of resting state connectivity between the insula and cingulate cortex. Hum Brain Mapp. 2009;30(9):2731–45.

120. Cavanna AE, Trimble MR. The precuneus: a review of its functional anatomy and behavioural correlates. Brain. 2006;129(3):564–83.

121. Tsolaki AC, Kosmidou V, Kompatsiaris IY, Papadaniil C, Hadjileontiadis L, Adam A, et al. Brain source localization of MMN and P300 ERPs in mild cognitive impairment and Alzheimer’s disease: a high-density EEG approach. Neurobiol Aging. 2017;55:190–201.

122. Papadaniil CD, Kosmidou VE, Tsolaki A, Tsolaki M, Kompatsiaris IY, Hadjileontiadis LJ. Cognitive 813MMN and P300 in mild cognitive impairment and Alzheimer’s disease: A high density EEG-3D vector 814field tomography approach. Brain Res. 2016;1648:425–33.

123. Jiang S, Qu C, Wang F, Liu Y, Qiao Z, Qiu X, et al. Using event-related potential P300 as an electrophysiological marker for differential diagnosis and to predict the progression of mild cognitive impairment: a meta-analysis. Neurol Sci. 2015;36(7):1105–12.

